# Comparative genome analysis of *Lactobacillus mudanjiangensis*, an understudied member of the *Lactobacillus plantarum* group

**DOI:** 10.1101/549451

**Authors:** Sander Wuyts, Camille Nina Allonsius, Stijn Wittouck, Sofie Thys, Bart Lievens, Stefan Weckx, Luc De Vuyst, Lebeer Sarah

**Affiliations:** University of Antwerp, Research Group Environmental Ecology and Applied Microbiology (ENdEMIC), Department of Bioscience Engineering, Antwerp, Belgium; Vrije Universiteit Brussel, Research Group of Industrial Microbiology and Food Biotechnology (IMDO), Faculty of Sciences and Bioengineering Sciences, Brussels, Belgium; University of Antwerp, Laboratory of Cell Biology and Histology, Antwerp Centre for Advanced Microscopy (ACAM), Antwerp, Belgium; KU Leuven, Laboratory for Process Microbial Ecology and Bioinspirational Management (PME&BIM), Department of Microbial and Molecular Systems (M2S), Campus De Nayer, Sint-Katelijne-Waver, Belgium

## Abstract

The genus *Lactobacillus* is known to be extremely diverse and consists of different phylogenetic groups that show a diversity roughly equal to the expected diversity of a typical bacterial genus. One of the most prominent phylogenetic groups within this genus is the *Lactobacillus plantarum* group which contains the understudied *Lactobacillus mudanjiangensis* species. Before this study, only one *L. mudanjiangensis* strain, DSM 28402^T^, was described but without whole-genome analysis. In this study, three strains classified as *L. mudanjiangensis*, were isolated from three different carrot juice fermentations and their whole-genome sequence was determined, together with the genome sequence of the type strain. The genomes of all four strains were compared with publicly available *L. plantarum* group genome sequences. This analysis showed that *L. mudanjiangensis* harbored the second largest genome size and gene count of the whole *L. plantarum* group. In addition, all members of this species showed the presence of a gene coding for a putative cellulose-degrading enzyme. Finally, three of the four *L. mudanjiangensis* strains studied showed the presence of pili on scanning electron microscopy (SEM) images, which were linked to conjugative gene regions, coded on plasmids in at least two of the strains studied.

**Author summary:** *Lactobacillus mudanjiangensis* is an understudied species within the *Lactobacillus plantarum* group. Since its first description, no other studies have reported its isolation. Here, we present the first four genome sequences of this species, which include the genome sequence of the type strain and three new *L. mudanjiangensis* strains isolated from fermented carrot juice. The genomes of all four strains were compared with publicly available *L. plantarum* group genome sequences. We found that this species harbored the second largest genome size and gene count of the whole *L. plantarum* group. Furthermore, we present the first scanning electron microscopy (SEM) images of *L. mudanjiangensis*, which showed the formation of pili in three strains that we linked to genes related to conjugation. Finally, we found the presence of a unique putative cellulose-degrading enzyme, opening the door for different industrial applications of these *Lactobacillus* strains.

## Introduction

The genus *Lactobacillus* is known to be extremely diverse [1]. Furthermore, it has been shown that different phylogenetic groups within this genus display a diversity roughly equal to the expected diversity of a typical bacterial genus [2–6]. Each of these phylogenetic groups can be recognized as an entity with unique properties and a distinct natural history, ecology, function and physiology [5]. Therefore, the study of these phylogenetic groups separately, as if they would be one genus, can be an interesting approach that might reveal new, previously overlooked, phylogenetic relationships and functional properties.

One of the more abundantly studied species within the genus *Lactobacillus*, is *Lactobacillus plantarum*. Previous genome-based phylogenetic studies have defined *L. plantarum* as a member of the *L. plantarum* group together with *Lactobacillus fabifermentans*, *Lactobacillus paraplantarum*, *Lactobacillus pentosus* and *Lactobacillus xiangfangensis* [1, 7]. In addition, the species *Lactobacillus herbarum* [8], *Lactobacillus plajomi* [9], *Lactobacillus modestisalitolerans* [9] and *Lactobacillus mudanjiangensis* [10] are closely related to *L. plantarum* and thus should be regarded as members of the *L. plantarum* group. *Lactobacillus mudanjiangensis* is a species that has been described for the first time in 2013 and that was isolated from a traditional pickle fermentation in the Heilongjiang province in China [10]. Since its first description, no other study has provided additional characterization or reported the isolation of other strains of the *L. mudanjiangensis* species. Therefore, currently, not a single genomic assembly of this species is publicly available. However, in in this study, four strains isolated from three different spontaneous carrot juice fermentations [11], were putatively classified as members of this *Lactobacillus* species.

Since the discovery of the mucus-binding pili, the fimbriae or adhesins, in *Lactobacillus rhamnosus* GG [12, 13], several comparative genomic studies have focused on exploring similar gene clusters in other lactobacilli, including the members of the *L. plantarum* group [1, 14–18]. Whereas these specific pili play an important role in cell surface adhesion, pili can be of importance for an array of other functions as well, ranging from biofilm formation to uptake of extracellular DNA via natural competence (type IV pili) or facilitation of DNA transfer via conjugation [19–21]. The latter is a process that uses conjugative pili to bring bacterial cells together and provide an interface to exchange macromolecules, such as DNA or DNA-protein complexes [20]. In general, such a conjugation system consists of three major components, namely *(i)* a relaxase (MOB) that will bind and knick the DNA at the origin of replication, *(ii)* a coupling protein (T4CP) that will couple the relaxase-DNA complex to *(iii)* a type IV secretion system (T4SS), which ultimately transfers the whole complex to the recipient cell [22–25]. Historically, these conjugation systems and their pili have been associated with conjugative plasmids only [23], one of the main drivers of horizontal gene transfer [22, 26]. However, recently, also integrative and conjugative elements (ICEs), which harbor conjugation systems as well, have been found to be another important driver of horizontal gene transfer [26–28].

This study aimed to provide more insights into the genomic features of the understudied *Lactobacillus mudanjiangensis* species, in relation to the other members of the *L. plantarum* group, using a comparative genomics approach. Therefore, the genome of the type strain of *L. mudanjiangensis* was sequenced together with three strains isolated from fermented carrot juice. These and other publicly available genome sequences were used to screen for *L. mudanjiangensis* species-specific properties, which included an analysis for the presence of genes related to pili formation and conjugation. In total, 304 genomes were subjected to an in-depth analysis focusing on the phylogenetic relationships as well as the predicted functional capacity of these strains.

## Results

The genome assembly of the type strain *L. mudanjiangensis* DSM 28402^T^ was analyzed together with the genome sequences of three putative *L. mudanjiangensis* strains isolated from carrot juice fermentations, namely AMBF197, AMBF209 and AMBF249, to confirm their putative classification as *L. mudanjiangensis* members. Furthermore, to allow comparison with other closely related *Lactobacillus* species and detection of *L. mudanjiangensis* species-specific properties, all publicly available genome sequences (NCBI Assembly database, 24/07/2018) of *L. plantarum* group members were included in this comparative genomics study, totaling the number of genomes analyzed to 304 (Table 1).

**Table 1.** An overview of the studied species and strains.

| Public data |  |  |  |
| --- | --- | --- | --- |
| Species | Number of genomes | Type strain | Reference |
| *L. fabifermentans* | 2 | DSM 21115<sup>T</sup> | [29] |
| *L. herbarum* | 1 | DSM 100358<sup>T</sup> | [8] |
| *L. mudanjiangensis* | 0 | DSM 28402<sup>T</sup> | [10] |
| *L. paraplantarum* | 5 | DSM 10667<sup>T</sup> | [30] |
| *L. pentosus* | 13 | DSM 20314<sup>T</sup> | [31] |
| *L. plantarum* | 278 | DSM 20174<sup>T</sup> | [32] |
| *L. xiangfangensis* | 1 | DSM 27103<sup>T</sup> | [33] |
| Type strains sequenced in this study |  |  |  |
| Species | Number of genomes | Type strain | Reference |
| *L. mudanjiangensis* | 1 | DSM 28402<sup>T</sup> | [10] |
| In-house isolates |  |  |  |
| Species | Number of genomes | Isolation source | Reference |
| *L. mudanjiangensis* | 3 | Spontaneously fermented carrot juice | [11] |

### Phylogeny of the *Lactobacillus plantarum* group

To obtain a detailed view on the phylogeny of *L. mudanjiangensis* in relationship to the whole *L. plantarum* group, a maximum likelihood phylogenetic tree was constructed, based on 612 single-copy core orthogroups, found with Orthofinder (Fig1 and S1 Fig). The resulting topology of this tree showed seven major clades, mostly following the species annotation as described in the NCBI Assembly database. However, these results exposed a few wrongly annotated genomic assemblies. For example, both *L. plantarum* MPL16 and *L. plantarum* AY01 were annotated as *L. plantarum* before, whereas here, they were found within a clade that contained the *L. paraplantarum* type strain. Similarly, *L. plantarum* EGD-AQ4 was found within the clade of the *L. pentosus* type strain, whereas it was annotated as *L. plantarum* before.

The type strain of *L. mudanjiangensis* formed a separate clade together with the strains AMBF197, AMBF209 and AMBF249 (Fig1). Based on its single-copy core orthogroups, this species was phylogenetically the most distant to *L. plantarum*, whereas its closest relative was *L. fabifermentans*, followed by *L. xiangfangensis*.

**Fig 1.**
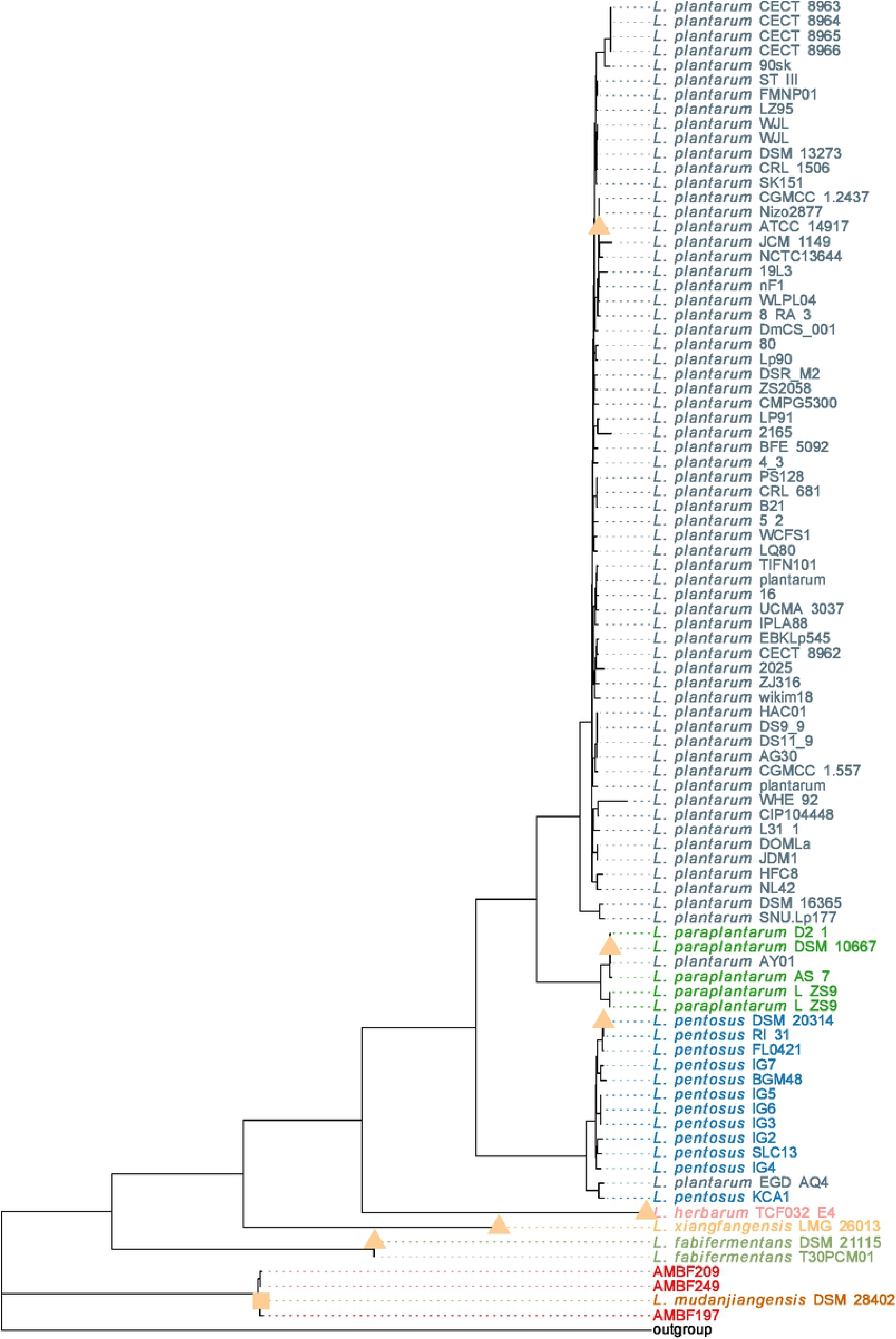
Maximum likelihood phylogenetic tree of the *Lactobacillus plantarum* group. The tree is based on the amino acid sequences of 612 single-copy marker genes. *Lactobacillus algidus* DSM 15638 was used as an outgroup. The tree was pruned to only keep 70 *L. plantarum* strains to avoid over-plotting. A complete tree can be found in the Appendix (S1 Fig). The branch length of the outgroup was shortened for better visualization. Each tip is colored, based on its species name as annotated in the NCBI Assembly database (where applicable), with dark blue for *L. plantarum*, light green for *Lactobacillus paraplantarum*, light blue for *Lactobacillus pentosus*, pink for *Lactobacillus herbarum*, light orange for *Lactobacillus xiangfangensis*, dark orange for *Lactobacillus mudanjiangensis* and red for the isolates obtained from the carrot juice fermentations. Type strains of each species are annotated with a triangle (NCBI) or a square (sequenced in this study).

### Low intraclade ANI values for *Lactobacillus pentosus* and *Lactobacillus plantarum*

To confirm that each major phylogenetic clade represented at least one different species, the pairwise average nucleotide identity (ANI) values of all genome assemblies were calculated (Fig2). Intraclade ANI values all exceeded the commonly used 95% species level threshold [34] for *L. mudanjiangensis* (99.0-99.4%), *L. fabifermentans* (99.7-99.9%) and *L. paraplantarum* (99.7-99.9%), whereas their interclade ANI values were far below this threshold, showing that these clades all represented a single species. However, this was not the case for *L. pentosus*, for which multiple pairwise comparisons led to intraclade ANI values below this threshold, suggesting that this phylogenetic clade contained at least two species (Fig2 and S2 Fig). This result was also found for some *L. plantarum* assemblies, although to a much lesser extent, compared to *L. pentosus*. Therefore, it was decided that, for subsequent analyses, *L. plantarum* was kept as one species, whereas *L. pentosus* was split into two groups, each representing one species. One species was designated as species *L. pentosus*, represented by a clade containing twelve genomic assemblies, including the type strain *L. pentosus* DSM 20314^T^. The other species, here referred to as clade 5a, was represented by two genomic assemblies (*L. pentosus* KCA1 and *L. plantarum* EGD-AQ4) (Figures 1 and S2 Fig). Finally, for *L. xiangfangensis* and *L. herbarum*, no intraclade comparisons could be performed, as only one genome assembly was available for these species.

**Fig 2.**
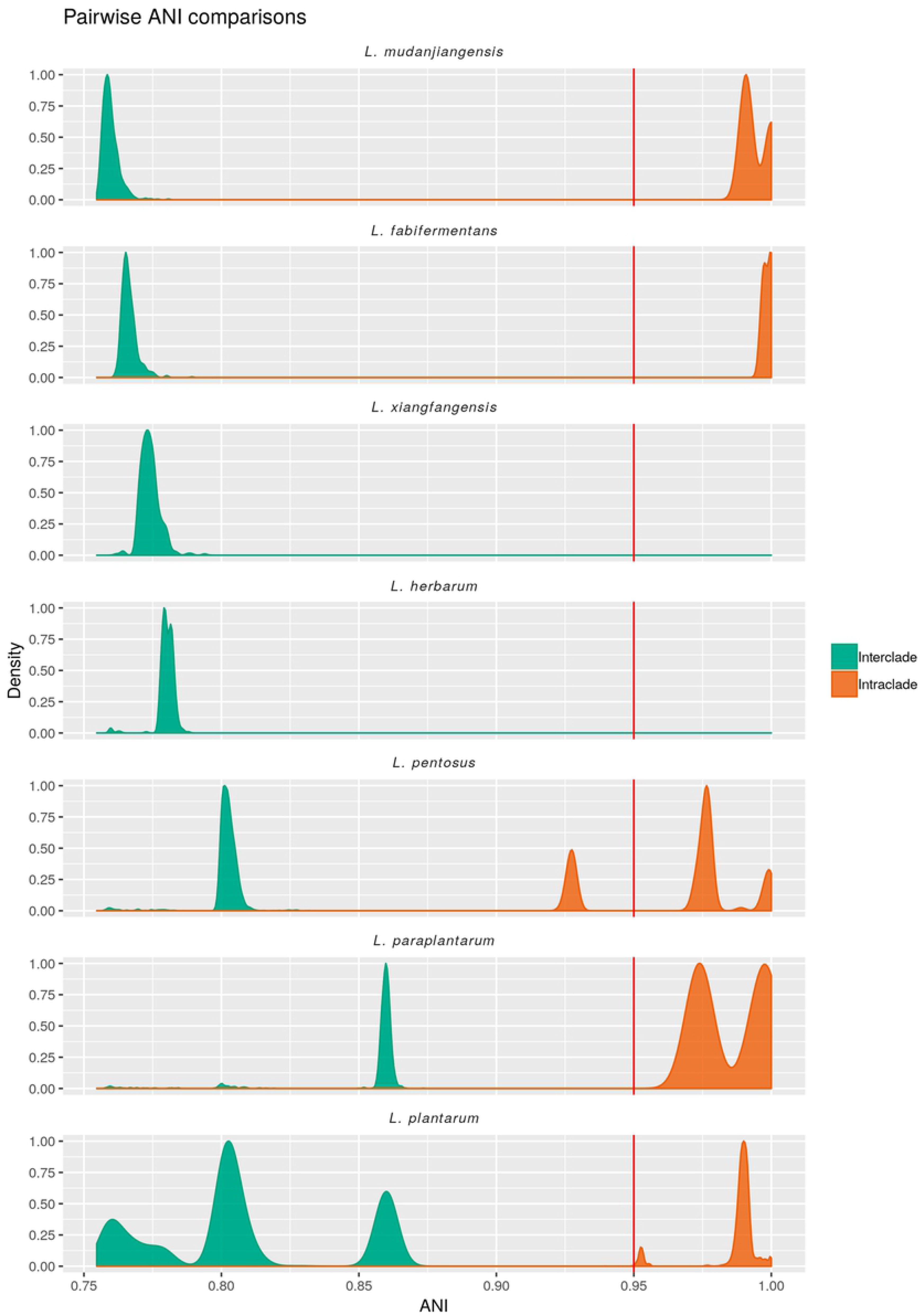
Density plot of all pairwise average nucleotide identity (ANI) comparisons for each *Lactobacillus plantarum* group species. In green all interclade comparisons are shown, whereas orange shows all intraclade comparisons. For *Lactobacillus xiangfangensis* and *Lactobacillus herbarum*, no intraclade comparisons could be performed, as only one genome assembly was available for these species.

### Genomic features of *Lactobacillus mudanjiangensis*

The above results confirmed the initial that strains AMBF197, AMBF209 and AMBF249 were members of the *L. mudanjiangensis* species. Therefore, here, the first four genomes of this species were presented. Their genome size varied between 3.4 Mb (strain DSM 28402^T^) and 3.6 Mb (strain AMBF209), whereas their GC content varied between 42.73% (strain AMBF209) and 43.06% (strain DSM 28402^T^) (Table 2). Finally, a high number of transfer RNA (tRNA) genes were found in all four strains.

**Table 2.** Genome characteristics of *Lactobacillus mudanjiangensis* strains.

| Genome | Total length (bp) | GC content (%) | Coding sequences | # tRNA genes |
| --- | --- | --- | --- | --- |
| AMBF197 | 3,501,388 | 42.85 | 3,463 | 65 |
| AMBF209 | 3,589,692 | 42.73 | 3,586 | 63 |
| AMBF249 | 3,554,025 | 42.83 | 3,503 | 71 |
| DSM 28402<sup>T</sup> | 3,389,962 | 43.06 | 3,346 | 66 |

A substantial difference in total genome length between the different species of the *L. plantarum* group was found (Fig3A). *Lactobacillus mudanjiangensis* showed a median estimated genome size of 3.53 Mb, the second largest of the whole *L. plantarum* group up to now. *Lactobacillus pentosus* showed the largest median estimated genome size (3.77 Mb), followed by *L. mudanjiangensis* (3.53 Mb) and *L. fabifermentans* (3.43 Mb), whereas for *L. xiangfangensis* (3.0 Mb) and *L. herbarum* (2.9 Mb) this size was much smaller. A remarkable high spread in genome length was found within strains belonging to the *L. plantarum* species, as their genome size ranged between 2.9 Mb and 3.8 Mb. Furthermore, *L. mudanjiangensis* showed a GC content of 42.9%, the lowest value within the whole *L. plantarum* group (Fig3B). Finally, regarding median gene count, similar trends were found as for the genome length, with *L. pentosus* showing the highest count, followed by *L. mudanjiangensis* and *L. fabifermentans*, whereas *L. xiangfangensis* and *L. herbarum* harbored the lowest numbers of genes (Fig3C).

**Fig 3.**
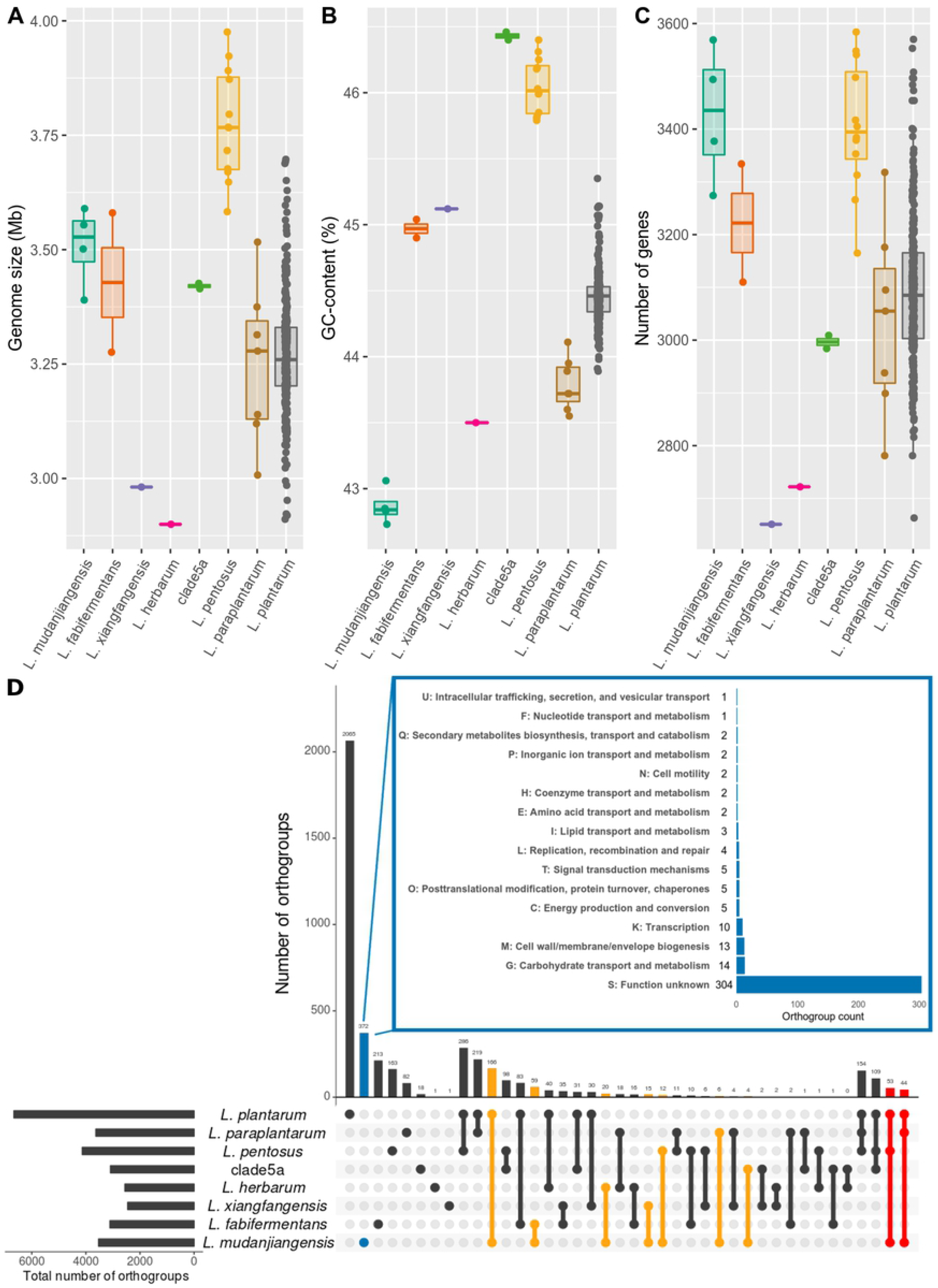
Estimated genome sizes, GC contents and gene counts of all genomes of *Lactobacillus plantarum* group species studied and predicted functional capacity of all unique *Lactobacillus mudanjiangensis* orthogroups. (A) Total genome size, (B) GC content and (C) gene counts of all genomes studied, colored by species. (D) Upset plot comparing shared orthogroup counts between the eight *L. plantarum* group species. Species-specific orthogroups for *L. mudanjiangensis* are colored in blue and the inset shows their functional category based on EggNOG classification. Uniquely shared orthogroups between *L. mudanjiangensis* and one other *L. plantarum* group member are colored in orange, whereas uniquely shared orthogroups between *L. mudanjiangensis* and two other species are colored in red.

In total, 947,588 genes were found in the whole *L. plantarum* group, with an average of 3,110 genes per genome. These genes were further clustered into 8,005 different orthogroups, leading to an average count of 2,924 orthogroups per genome. The differences between these numbers was due to the fact that some genes were found in multiple copies within one genome, which clustered together in a single orthogroup. Of all these orthogroups, 2,172 were defined as core orthogroups and 5,833 as accessory orthogroups (see Sections 1.2.2 and 4.2.3 for the definition of these terms). A detailed overview of the number of genes and core and accessory orthogroups can be found in S2 Table. Subsequently, the distribution of orthogroups between the different *L. plantarum* group members was explored (Fig3D). The species with the highest number of species-specific orthogroups was *L. plantarum*. With 2,065 species-specific orthogroups, it greatly outnumbered all other species, although this number was most probably biased, due to the higher number of sequenced genomes available for *L. plantarum* compared with the other *L. plantarum* group species. It was followed by *L. mudanjiangensis* (Fig3D, blue) that contained 372 species-specific orthogroups and *L. fabifermentans* harboring 213 species-specific orthogroups. Furthermore, *L. plantarum* and *L. pentosus* shared the highest number of uniquely shared orthogroups (286), followed by the combination of *L. plantarum* and *L. paraplantarum* (219 uniquely shared orthogroups), which seemed to be in line with the phylogeny described in Fig1. In contrast, *L. mudanjiangensis* shared more unique orthogroups with the phylogenetically distant *L. plantarum* (166 orthogroups; Fig3D, yellow) than it did with the most closely related species, *L. fabifermentans* (59 orthogroups).

To get more insights into the unique properties of *L. mudanjiangensis*, all 372 species-specific orthogroups were further classified using the EggNOG database (inset Fig3D). However, this resulted in a vast majority of orthogroups (304), classified under ‘Category S: Function unkown’, showing that further experimental work on functional gene validation is necessary. Besides this, most orthogroups belonged to category G (carbohydrate transport and metabolism, 14 orthogroups), followed by category M (cell wall/membrane/envelope biogenesis, 13 orthogroups).

### *Lactobacillus mudanjiangensis* harbors a potential cellulose-degrading enzyme

Carbohydrate transport and metabolism (category G) was found to be the most abundantly characterized category among the *L. mudanjiangensis* species-specific orthogroups. Further examination of the 14 unique orthogroups that were detected in this category, revealed the presence of a gene in all four strains annotated as endoglucanase E1, which is involved in the conversion of cellulose polymers into simple saccharides [35]. A BLAST search of the DNA sequences of this gene to the NCBI nt database showed a best scoring hit (26% coverage and 69% identity) with a member of the *Herbinix* species (GenBank accession number LN879430). The *Herbinix* genus contains cellulose-degrading bacteria [36]. This result also showed that this gene was not found in any other member of the *Lactobacillus* Genus Complex (LGC), or any other LAB, and confirmed its uniqueness to *L. mudanjiangensis*. Since endoglucanases are classified as glycosyl hydrolases (GHs), GHs were predicted for all genomes. Indeed, for all four *L. mudanjiangensis* strains, this endoglucanase E1 gene was classified as belonging to the GH5 1 family, which was a family uniquely found in *L. mudanjiangensis*. Although this GH family showed some degree of polyspecificity, the majority of enzymes (22 of 24 enzymes characterized) are reported as endoglucanases [37]. Together, these results thus pointed towards the presence of a novel putative cellulose-degrading enzyme in all four *L. mudanjiangensis* strains.

### Presence of a putative conjugative system in *Lactobacillus mudanjiangensis*

The second most abundant category of *L. mudanjiangensis*-specific orthogroups, excluding category S (function unknown), were genes related to ‘cell wall, membrane, or envelope biogenesis’ (category M). Examination of the annotation of the genes belonging to these orthogroups did not reveal any new insights, as many of them were annotated as hypothetical proteins. Therefore, scanning electron microscopy (SEM) was performed to screen the cell surfaces of these four strains in more detail. This analysis revealed that three of the four strains (*L. mudanjiangensis* DSM28402^T^, AMBF209 and AMBF249) formed pili or fimbriae, connecting different cells to each other as well as cells to an undefined structure (Fig4A).

**Fig 4.**
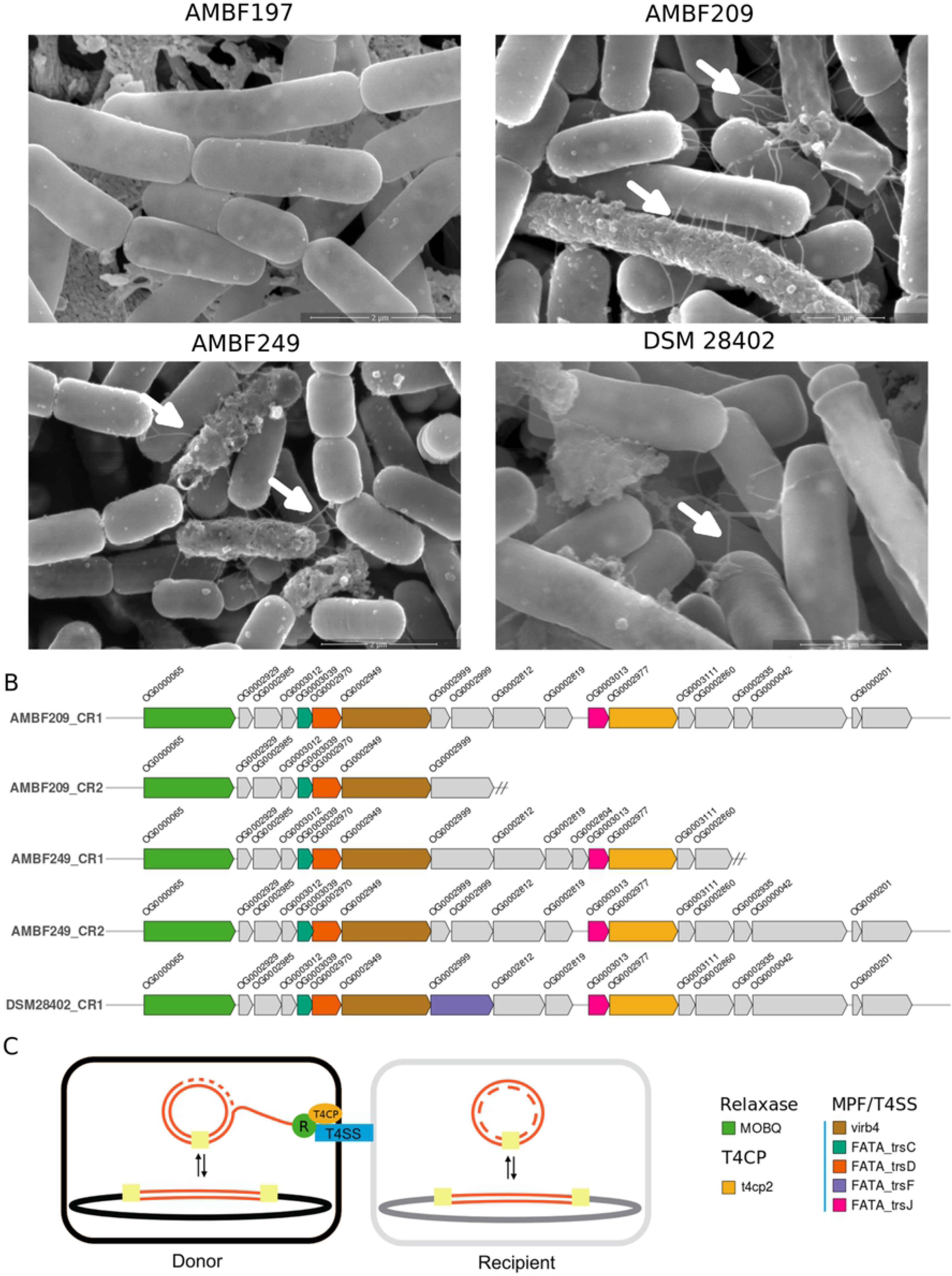
Scanning electron microscopy (SEM) and genes related to conjugation. (A) SEM images of all four *Lactobacillus mudanjiangensis* strains studied. White arrows indicate putative conjugative pili. (B) Gene clusters encoding a putative conjugation system, colored by their potential function, as classified by CONJSCAN. The text above each gene shows its matching orthogroup. (C) Schematic model representing the process of bacterial conjugation with all three mandatory elements. Scheme adapted from [23]. (R, relaxase; T4CP, Type IV coupling protein; T4SS, Type IV secretion system).

To identify the genes encoding these pili, all genome sequences of *L. mudanjiangensis* were screened for the presence of genes associated with these kinds of phenotypes. These included the *spaCBA* gene cluster, which has been linked with probiotic properties in *L. rhamnosus*, due to better adhesion to intestinal epithelial cells [38, 39], as well as secretion systems based on pili, such as the type II and type IV secretion systems [25, 40]. In this study, no *spaCBA* gene cluster was found. However, further exploration revealed the presence of a conjugation system in at least three of the four *L. mudanjiangensis* strains examined (AMBF209, AMBF249 and DSM 28402^T^).

Two complete conjugation systems containing all three mandatory parts (Fig4C) were found in *L. mudanjiangensis* AMBF209 and AMBF249, whereas one complete conjugation system was found in *L. mudanjiangensis* DSM 28402^T^ (Fig4B). For all four *L. mudanjiangensis* strains, the relaxase gene of this conjugation system was classified as a member of the MOBQ class, whereas the coupling protein was classified as a T4CP2. The MPF system, which harbored the putative pilus, was further classified as belonging to the class MPFFATA, which groups the MPF systems of Gram-positive bacteria [24]. VirB4 was identified as the ATPase motor of this MPF system. Furthermore, this MPF system contained three accessory genes (*trsC*, *trsD* and *trsJ)* in *L. mudanjiangensis* AMBF209 and AMBF249, whereas four accessory genes were annotated in *L. mudanjiangensis* DSM 28402^T^ (*trsC*, *trsD*, *trsF* and *trsJ)* (Fig4B). Homologs for the genes *trsC* and *trsD* were already previously identified, with *trsC* coding for a VirB3 homolog, which is linked to the formation of the membrane pore, and *trsD* coding for another homolog of VirB4, the conjugation ATPase [24]. In contrast, both *trsF* and *trsJ* are poorly characterized.

Further analysis of the genes surrounding the annotated conjugation genes showed that this genomic region contained 18 to 19 open reading frames, most of them annotated as hypothetical proteins (Fig4B and S3 Table). However, a bacteriophage peptidoglycan hydrolase domain was found in orthogroup OG0002812 in both *L. mudanjiangensis* AMBF209 conjugation region 1 (AMBF209 CR1) and *L. mudanjiangensis* AMBF249 conjugation region 2 (AMBF249 CR2), making it a VirB1-like protein [41]. In *Agrobacterium tumefaciens*, the VirB1 protein provides localized lysis of the peptidoglycan cell wall to allow insertion of the T4SS [42]. A similar domain, also known to harbor peptidoglycan lytic activity, was found in *L. mudanjiangensis* DSM28402 CR1 (orthogroup OG0002812). Finally, another conserved domain was found in all five gene regions clustered in orthogroup OG0003012, annotated as T4SS CagC, which was shown to be a VirB2 homolog. VirB2 is the major pilus component of the type IV secretion system of *A. tumefaciens*, which is the main building block for extension and retraction of the conjugative pilus [43–45]. Taken together, these results showed the presence of pili in three *L. mudanjiangensis* strains (AMBF209, AMBF249 and DSM 28402^T^), which after genomic analysis were hypothesized to be part of a conjugation system.

Finally, genome analysis of all other *L. plantarum* group members showed that, in contrast to an initial belief, the presence of a complete conjugation system was not unique to *L. mudanjiangensis* (S1 Table). All three necessary genes were also found in 58 of 275 *L. plantarum* strains, two of seven *L. paraplantarum* strains and four of twelve *L. pentosus* strains. In contrast, the system was completely absent in clade5a, *L. herbarum*, *L. xiangfangensis* and *L. fabifermentans*.

### Plasmid reconstruction from genome data

Many conjugation systems are coded on plasmids [27]. Therefore, all four *L. mudanjiangensis* genomes were screened for plasmid presence using Recycler [46]. Plasmids were only found in two of four genome assemblies, namely *L. mudanjiangensis* AMBF209 and AMBF249 (Fig5). Both strains harbored a plasmid of 27.3 Kb with 33 predicted genes, which after pairwise alignment of the plasmid were foun to be exactly the same. Subsequently, the presence of a conjugation system on these plasmids was confirmed using CONJScan. Further examination showed that the plasmid exactly matched the above described AMBF209 CR2 and AMBF249 CR1 gene regions (Fig4B). Regarding annotated genes, 13 of 33 gene products were predicted as hypothetical proteins by Prokka. Further annotation using the EggNOG database revealed that most genes were mapped to category S (function unknown), followed by category L (replication, recombination and repair) and category C (energy production and conversion). Finally, a BLAST search was performed against the NCBI nt database to explore whether a similar plasmid was already described in the literature. This resulted in a best matching hit, showing 97% identity and 76% query coverage with plasmid pKLC4 of *Leuconostoc carnosum* JB16 (GenBank accession number CP003855). This plasmid was found in a strain isolated from kimchi and has a length of 36.6 Kb [47], indicating that some deletions occurred in the *L. mudanjiangensis* plasmids. All together, these results showed that *L. mudanjiangensis* strains AMBF209 and AMBF249 carried the same conjugative plasmid, of which the encoded gene functions are poorly characterized.

**Fig 5.**
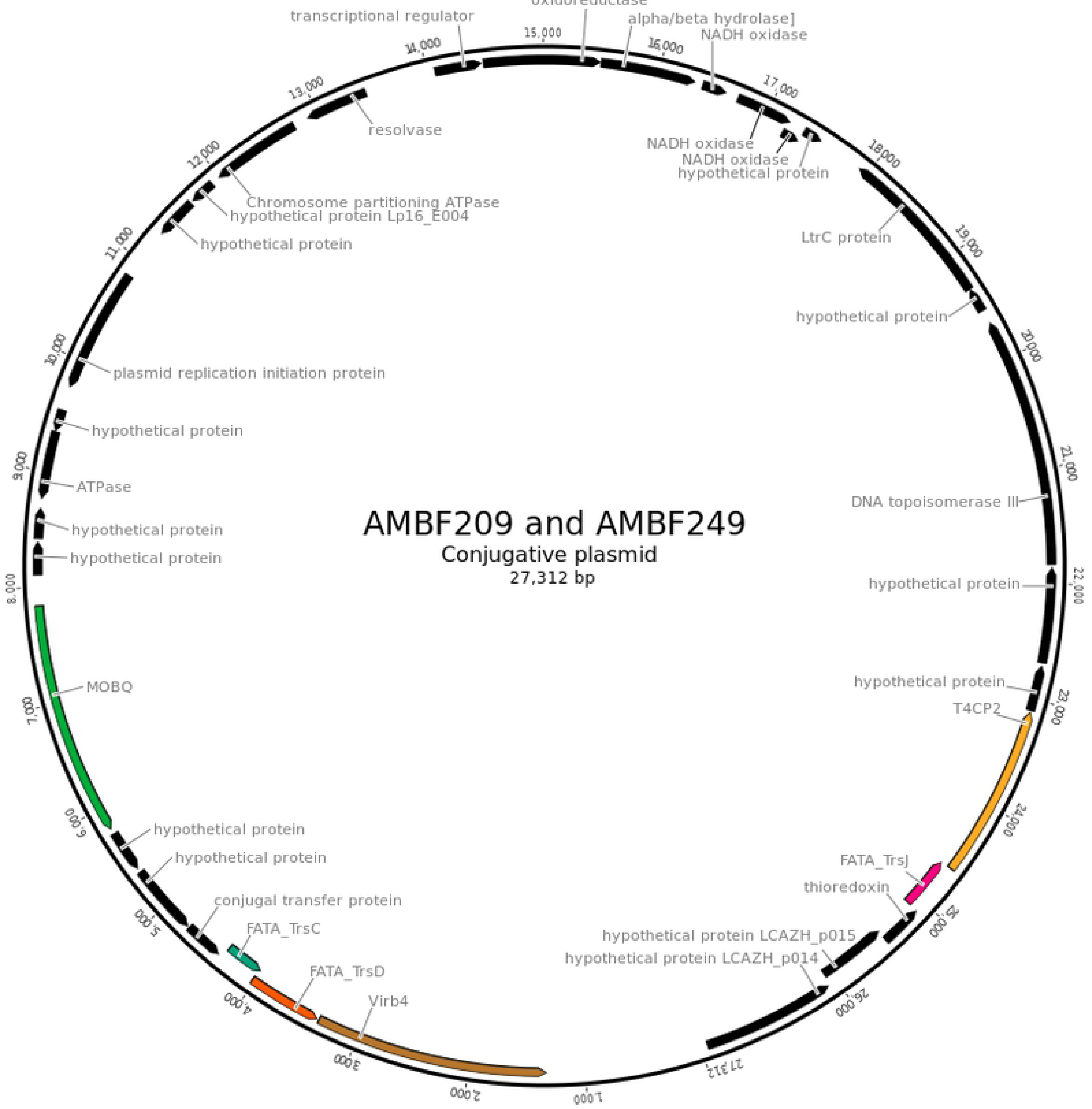
Plasmid map of the predicted conjugative plasmid of *Lactobacillus mudanjiangensis* AMBF209 and AMBF249. Genes are colored according to their annotation, as defined in Fig4.

Since only two of five conjugation regions (Fig4) were plasmid-encoded, an additional analysis was performed to assess whether the other three conjugation systems could be part of an ICE. For this, all four *L. mudanjiangensis* genomes were analyzed similar to a recently published method [26]. However, ICE regions usually contain repeats, such as transposases, leading to fragmentation of these ICE regions, if short read sequencing technology is used [26]. Therefore, these analysis methods usually require a complete genome for proper ICE identification. The assembly state of the four genomes made it thus hard to correctly interpret the results obtained.

## Discussion

In this study, the genome sequence of the *L. mudanjiangensis* type strain DSM 28402^T^ was presented together with the genomes of three new *L. mudanjiangensis* strains, AMBF197, AMBF209 and AMBF249, which were isolated from three different spontaneous carrot juice fermentations [11]. Since previous phylogenetic analysis of this species, using the 16S rRNA, *pheS*, and *rpoA* genes, showed a close genetic relatedness with the members of the *L. plantarum* group [10], it was decided to study these genomes in relation to the closely related members of the *L. plantarum* group with a comparative genomics approach. A maximum likelihood phylogenetic tree confirmed that *L. mudanjiangensis* was closely related to all other *L. plantarum* group members. Furthermore, pairwise ANI analysis confirmed that the three strains isolated from carrot juice fermentations were indeed members of the *L. mudanjiangensis* species. Yet, this analysis also revealed that several genomes annotated as *L. pentosus* and *L. plantarum* showed intraclade ANI values below the commonly used cut-off value of 95% identity [34]. Especially for *L. pentosus*, low intraclade ANI values (a maximum of 92,3%) were found, meaning that if the 95% cut-off is strictly applied, two genome assemblies could not be seen as members of the *L. pentosus* species. One of them represented the genome sequence of the vaginal isolate *L. pentosus* KCA1 [48], the other a genome sequence that has been annotated as *L. plantarum* EGD-AQ4, isolated from a fermented bamboo shoot product [49]. The fact that these two genomes showed lower intraclade ANI values could mean that these bacteria are undergoing a speciation process or that this strict ANI value cut-off should be revised, as it potentially separates members from the same species into different species.

The estimated genome size of *L. mudanjiangensis* was the second largest of the whole *L. plantarum* group found up to now. The same trend was found for the gene count per species. This means that *L. mudanjiangensis* harbored one of the largest genomes of the whole LGC, since *L. plantarum* and especially *L. pentosus* are known to be among the lactobacilli with the largest genome size and gene counts [5, 50, 51]. For *L. plantarum*, this large genome and also its pangenome size have been coupled to a nomadic lifestyle [5, 50]. This lifestyle comes with a high genetic diversity, which is associated to the possibility to survive and thrive in many different ecosystems. This could possibly also be applied to *L. mudanjiangensis*, supported by the fact that *L. mudanjiangensis* showed a slightly larger number of GHs than *L. plantarum* (Data not shown), indicating that this species is capable of transforming and metabolizing a broad spectrum of carbohydrate sources. This observation was confirmed by the fact that the type strain was found to be capable of producing organic acids from at least 21 different carbon sources [10]. Furthermore, here, a putative cellulose-degrading enzyme, annotated as endoglucanase E1, was found in all four *L. mudanjiangensis* strains, which was not found in any other LGC genome so far. Moreover, cellulose-degrading enzymes were never found in LAB up to now. Cellulose is the most abundant organic polymer on earth [52], the most important skeletal component in plants in general [52] and the most abundant crude fiber in carrots [53]. *Lactobacillus mudanjiangensis’* putative capability to degrade this fiber into glucose might allow members of this species to survive in many different plant-related ecosystems. Fermented carrot juices and fermented pickles are examples of such ecosystems, where three and one of the strains studied were isolated from, respectively. Together, these results suggested that, similar to *L. plantarum*, a nomadic or otherwise a plant-adapted lifestyle could be assigned to *L. mudanjiangesis*.

SEM analysis revealed the presence of pili or fimbriae in three of the four *L. mudanjiangensis* strains studied. In this study, the observation of pili in *L. mudanjiangensis* was associated with bacterial conjugation. Three of the four strains were found to carry at least one complete putative conjugation region, including a gene that possibly codes for a VirB2 homolog, the major subunit of a conjugation-related pilus [43–45]. The three strains that harbored this conjugation region all showed pili formation on the SEM images, whereas this was not the case for strain AMBF197, which lacked this region. These results suggested that the detected pili might play a role in cell to cell contact during the conjugation process, although this was not yet experimentally validated.

Conjugation is one of the main drivers of horizontal gene transfer and is commonly associated with conjugative plasmids [22, 26]. Here, two of the five conjugative regions found were plasmid-associated and the two plasmids found were exactly the same for both *L. mudanjiangensis* AMBF209 and AMBF249, although these strains were isolated from different household carrot juice fermentations [11]. Previous studies also identified and described conjugative plasmids in other *Lactobacillus* species, such as *Lactobacillus brevis* [54], *Lactobacillus casei* [55], *Lactobacillus gasseri* [56], *Lactobacillus hokkaidonensis* [57], *L. plantarum* [58] and *Lactobacillus reuteri* [59]. Genes on these plasmids often code for proteins involved in detoxification, virulence, antibiotic resistance and ecological interactions [22], which could give them a fitness advantage in certain environments. Here, apart from the conjugation-related genes, many genes were annotated as hypothetical proteins on the conjugative plasmid. However, since this plasmid showed great similarity with a plasmid from a *Leuconostoc* strain, which was isolated from fermented kimchi [47], it could potentially harbor genes that are beneficial for survival on plants or in a fermented vegetable environment.

In conclusion, in this study, the genome sequences of four *L. mudanjiangensis* strains were studied in relation to the closely related members of the *L. plantarum* group. Comparative genome analysis of this phylogenetic group found two wrongly annotated genome assemblies and intraclade ANI values below the commonly used species delimitation threshold for *L. plantarum* and *L. pentosus*. Furthermore, *L. mudanjiangensis* harbored one of the largest genomes and the highest gene counts of the 1. *L. plantarum* group. Together with its broad repertoire of GHs and its potential capability to degrade cellulose, either a nomadic or plant-adapted lifestyle could be assigned to *L. mudanjiangensis*. Finally, three of the four *L. mudanjiangensis* strains studied showed the presence of pili on SEM images, which were linked to conjugative gene regions. For two strains, *L. mudanjiangensis* AMBF209 and AMBF249, these regions were plasmid-associated. Further experimental studies, such as phenotypic growth curve-based screenings, conjugation experiments and the creation of knock-out mutants, are necessary to characterize the plasmid found and to confirm the link between the pili observed and this conjugation gene region.

## Materials and methods

### Sequencing of the *Lactobacillus mudanjiangensis* type strain and downloading of publicly available assemblies

The type strain of *L. mudanjiangensis* [*L. mudanjiangensis* DSM 28402^T^ (= LMG 27194^T^ = CCUG 62991^T^)] was purchased from a public microorganism collection (BCCM-LMG, Ghent, Belgium). The strain was grown overnight in de Man-Rogosa-Sharpe (MRS) medium (Carl Roth, Karlsruhe, Germany) and DNA was extracted using the NucleoSpin 96 tissue kit (Macherey-Nagel, Düren, Germany), with an extra cell lysis step using 20 mg/mL of lysozyme (Sigma-Aldrich, St. Louis, MO, USA) and 100 U/mL of mutanolysin (Sigma-Aldrich). Whole-genome sequencing was performed using the Nextera XT DNA Sample Preparation kit (Illumina, San Diego, CA, USA) and the Illumina MiSeq platform, using 2 × 250 cycles, at the Laboratory of Medical Microbiology (University of Antwerp, Antwerp, Belgium) in the case of the strains AMBF197, AMBF249 and DSM28402^T^ or 2 × 300 cycles at the Center of Medical Genetics Antwerp of the University of Antwerp for strain AMBF209. Assembly of the genome sequence was performed using SPAdes v 3.12.0 [60]. In addition, all genome sequences annotated as *L. fabifermentans*, *L. herbarum*, *L. paraplantarum*, *L. pentosus*, *L. plantarum* and *L. xiangfangensis* were downloaded from the National Center for Biotechnology Information (NCBI) Assembly database on 24/07/2018, using in-house scripts. In total, 310 genomes were used as an input for quality control.

### Quality control and annotation

Basic genome characteristics, including genome size, GC content and the N50 value, were estimated using Quast 4.6.3 [61]. The quality of the genome assemblies was evaluated using the Quast output. After visualization of several quality control parameters using ggplot2 [62], genomes with a N50 value *<* 25,000 bp and a number of undefined nucleotides (N) per 100,000 bases *>* 500 were discarded. An overview of all genome sequences and strains that passed this quality control (304 assemblies) can be found in S1 Table. Finally, Prokka 1.12 [63] was used to predict and annotate genes for all genome sequences. In addition to its internal databases, a customized genus-specific BLAST database was used for higher quality annotation with Prokka’s *–usegenus* option. This database was created using BLAST [64, 65] and all complete *Lactobacillus* genomes found in the NCBI Assembly database. Genes encoding glycosyl hydrolases (GHs) were detected by scanning all genomes against hidden Markov model (HMM) profiles of the CAZyme families [66]. The profiles were downloaded from the dbCAN webserver [67] and queried using HMMSCAN [68]. An E-value of 1 × 10-15 and a coverage of 0.35 were used as cut-off, similar to what has been described before [69].

### Defining the pangenomes of all *Lactobacillus plantarum* group species

To define the pangenome, all genes were clustered into orthogroups using OrthoFinder [70] and further analyzed in R [71]. Here, a core orthogroup is defined as an orthogroup present in more than 95% of a set of genomes. All other orthogroups are defined as accessory orthogroups. An upset plot was created using the R package UpSetR [72]. Unique orthogroups belonging to *L. mudanjiangensis* were further annotated using EggNOG-mapper [73] and visualized using ggplot2 [62].

### Phylogenetic tree construction

Single-copy core orthogroups found by Orthofinder were used as input for the construction of a phylogenetic tree. *Lactobacillus algidus* DSM 15638 (NCBI Assembly accession number GCA 001434695) served as an outgroup, as it is the species most closely related to the *L. plantarum* group [1]. The first protein sequence of each fasta file of the single-copy core orthogroups was compared with a BLAST database of all genome proteins of the outgroup’s genome sequence. All hits with a coverage *>* 75% and a percentage similarity *>* 50% were added to the alignment of each orthogroup. These alignments, on amino acid level, were concatenated into a supermatrix that was used in RaxML 8.2.9 [74], to build a maximum likelihood phylogenetic tree with the *–a* option, which combines a rapid bootstrap algorithm with an extensive search of the tree space, starting from multiple different starting trees. The tree and subtrees were plotted with the R package ggtree [75].

### Average nucleotide identity

All pairwise ANI values were calculated with the Python pyani package [76], using a BLASTN approach [64, 65] based on the methodology described by Goris *et al.* [77].

### Scanning electron microscopy

To assess the presence or absence of pili or fimbriae on the cell surface of *L. mudanjiangensis* strains AMBF197, AMBF209, AMBF249 and DSM 28402^T^, SEM was performed. To this end, the bacterial strains were grown overnight (MRS medium, 37°C), gently washed with phosphate-buffered saline (per liter: 56 g of NaCl, 1.4 g of KCl, 10.48 g of Na2HPO4, 1.68 g of KH2PO4; pH 7.4) and spotted on a gold-coated membrane [(approximately 5 × 107 colony forming units (CFU) per membrane]. Bacterial spots were fixed with 2.5% (m/v) glutaraldehyde in 0.1 M sodium cacodylate buffer (2.5% glutaraldehyde, 0.1 M sodium cacodylate, 0.05% CaCl2.2H2O; pH 7.4) by gently shaking the membrane for 1 h at room temperature, followed by a further overnight fixation at 4 °C. After fixation, the membranes were washed three times for 20 min with cacodylate buffer (containing 7.5% [m/v] saccharose). Subsequently, the bacteria were dehydrated in an ascending series of ethanol (50%, 70%, 90% and 95%, each for 30 min at room temperature and 100% for 2 × 1 h and 1 × 30 min) and dried in a Leica EM CPD030 (Leica Microsystems Belgium, Diegem, Belgium). The membranes were mounted on a stub and coated with 5 nm of carbon (Leica Microsystems Belgium) in a Leica EM Ace 600 coater (Leica Microsystems Belgium). SEM imaging was performed using a Quanta FEG250 SEM system (Thermo Fisher, Asse, Belgium) at the Antwerp Centre for Advanced Microscopy (ACAM, University of Antwerp) and Electron Microscopy for Material Science group (EMAT, University of Antwerp).

### Detection of genomic clusters encoding pili or fimbriae

To screen for the presence of the *spaCBA* gene cluster, the gene cluster that is responsible for expression of the fimbriae in *L. rhamnosus* GG [38, 39], a BLAST search [64, 65] on protein level was performed against a BLAST database constructed for each genome separately. The gene sequences of *spaA* (NCBI GenBank accession number BAI40953.1), *spaB* (BAI40954.1) and *spaC* (BAI40955.1) were used as queries. Furthermore, the genomes were screened for genes encoding pili-related protein secretion systems, using the predicted amino acid sequences as query and the TXSScan definitions and profile models [25] as references in MacSyFinder v1.0.5 [78]. As only genes related to conjugation systems were found, all protein sequences of all genomes were again scanned, this time using the CONJScan definitions and profile models [23, 26] using MacSyFinder. In brief, a conjugation region was only considered if the conjugation genes were separated by less than 31 genes, except for genes encoding relaxases that can be separated by maximal 60 genes. The region was considered conjugative when it contained genes coding for *(i)* a VirB4/TraU homolog, *(ii)* a relaxase, *(iii)* a type 4 coupling protein (T4CP) and *(iv)* a minimum number of mating-pair formation (MPF) type-specific genes [26]. For both scans, hits with alignments covering *>* 50% of the protein profile and with an independent E-value ¡ 10^*−*3^ were kept for further analysis (default parameters) in R [71]. Conserved domain analysis of genes of interest was performed using the NCBI Conserved Domain web interface [79]. The gene regions were visualized using the R package gggenes (available at https://github.com/wilkox/gggenes).

### Plasmid identification

Detection and reconstruction of plasmids in the different *L. mudanjiangensis* strains was performed using Recycler v0.7 [46], with the original fastq files and SPAdes assembly graphs as input. The assembled plasmids were annotated with Prokka and further characterized by scanning against the EggNOG database, as described in above. The presence of a conjugation system was confirmed with CONJScan, as described above. The percentage identity between the different plasmids found was assessed using BLAST [64, 65]. The similarity with any previously described plasmid was checked by performing a BLAST search [64, 65] against the NCBI nucleotide (nt) database. A plasmid map was created using Geneious v8 [80].

### Delimitation of integrative and conjugative elements

The presence of ICEs was explored by a similar approach as the pipeline described previously [26]. Briefly, all strict core genes, *i.e.* genes present in all strains of *L. mudanjiangensis* were found using the Orthofinder output (see above). Next, all flanking core genes of each conjugative region were identified. Since within one species an ICE is expected to be found between the same core orthogroups, the flanking core genes of each conjugative region found were evaluated to determine whether or not it could be defined as an ICE.

### Accession number(s) and data availability

Sequencing data and genome assemblies are available at the European Nucleotide Archive under the accession number ERP111972. The complete pipeline can be found on GitHub (https://github.com/swuyts/mudAnalysis).

## Acknowledgments

We thank Eline Oerlemans, Ilke De Boeck, Ines Tuyaerts, and all other members of the ENdEMIC group for their general assistance and/or fruitful discussions. In addition, we also thank Charlotte Claes and Arvid Suls (Centre of Medical Genetics, University of Antwerp, Antwerp, Belgium) for their valuable input regarding whole-genome sequencing. Furthermore, we would like to thank the Antwerp Centre for Advanced Microscopy (ACAM, University of Antwerp) for processing and imaging of the SEM samples and the Electron Microscopy for Material Science group (EMAT, University of Antwerp) for the use of the environmental SEM (Quanta 250 FEG).

## Supporting information

**S1 Fig. Maximum likelihood phylogenetic tree of the whole *Lactobacillus plantarum* group constructed using 304 genome assemblies and the amino acid sequences of their 612 single-copy marker genes with *Lactobacillus algidus* DSM 15638 as outgroup.** The branch length of the outgroup was shortened for better visualization. The colors represent the species as annotated in the National Center for Biotechnology Information (NCBI) Assembly database. Type strains of each species are annotated with a triangle (NCBI) or a square (sequenced in-house).

**S2 Fig. Alternative vizualisation of all pairwise average nucleotide identity (ANI) comparisons for each *Lactobacillus plantarum* group species, as defined by the clades in Fig1.** In green all inter-clade comparisons are shown, while orange shows all intra-clade comparisons. For *Lactobacillus xiangfangensis* and *Lactobacillus herbarum*, no intra-clade comparisons could be performed, as only one genomic assembly was available for these species.

**S1 Table. Overview of all of the publicly available *Lactobacillus plantarum* group genomes used in this study.** If available, the strain name is shown in the second column. The clade name is in accordance to the phylogenetic tree of Fig1. The last column indicates whether a conjugation system was found or not.

S2 Table. Number of genomes, core orthogroups, accessory orthogroups, average orthogroups per genome and average genes per genome of the *Lactobacillus plantarum* group as a whole and all of its members separately.

**S3 Table. An overview of all genes found in the *Lactobacillus mudanjiangensis* conjugation operons.** The CONJscan column contains the annotation resulting from running the CONJscan tool.

